# New factors for protein transport identified by a genome-wide CRISPRi screen in mammalian cells

**DOI:** 10.1101/556746

**Authors:** Laia Bassaganyas, Stephanie J. Popa, Max Horlbeck, Anupama Ashok, Sarah E. Stewart, Cristian M. Butnaru, Nathalie Brouwers, Kartoosh Heydari, Jean Ripoche, Jonathan Weissman, Randy Schekman, Vivek Malhotra, Kevin Moreau, Julien Villeneuve

## Abstract

Protein and membrane trafficking pathways are critical for cell and tissue homeostasis. Traditional genetic and biochemical approaches have shed light on basic principles underlying these processes. However, the list of factors required for secretory pathways function remains incomplete, and mechanisms involved in their adaptation poorly understood. Here, we present a powerful strategy based on a pooled genome-wide CRISPRi screen that allowed the identification of new factors involved in protein transport. Two newly identified factors, TTC17 and CCDC157, localized along the secretory pathway and were found to interact with resident proteins of ER-Golgi membranes. In addition, we uncovered that upon TTC17 knockdown, the polarized organization of Golgi cisternae was altered, creating glycosylation defects, and that CCDC157 is an important factor for the fusion of transport carriers to the Golgi complex. In conclusion, our work identified and characterized new actors in the mechanisms of protein transport and secretion, and opens stimulating perspectives for the use of our platform in physiological and pathological contexts.

## Introduction

The molecular machinery underlying the processes of protein transport and secretion has been conserved from yeast to mammalian cells, being essential to maintain specificity and communication between organelles, exocytosis and endocytosis. Fully one-third of proteins navigate the secretory pathway, entering this system via the endoplasmic reticulum (ER). They are then transported via cellular compartments, including the Golgi apparatus and transport carriers, to be targeted to their final destination (Lee et al., 2004). Beyond this apparent simple view, processes underlying protein transport and secretion are gaining continuous complexity with recent demonstrations that cell compartments establish cross regulatory mechanisms with numerous membrane contact sites (Wu et al., 2018), are endowed of tightly regulated dynamics (Valm et al., 2017), and stand at the crossroad of signaling pathways where inputs and outputs are integrated and coordinated (Luini and Parashuraman, 2016).

Yeast genetic studies and *in vitro* biochemical approaches have initially been used to discover the basal protein transport machinery (Novick et al., 1980; Braell et al., 1984). More recently, arrayed RNA interference screens at the genome scale or targeting kinases/phosphatases, expanded the list of functional and regulatory components of secretory pathways in metazoan cells (Asensio et al., 2010; Bard et al., 2006; Chia et al., 2012; Simpson et al., 2012; Wendler et al., 2010). In these studies, artificial secreted reporters such as horseradish peroxidase (HRP) or firefly luciferase preceded by a signal sequence (ss) were used for detection. Alternatively, Golgi membrane organization or transport to the cell surface of fluorescently labeled exogenous transmembrane proteins such as Vesicular Stomatitis Virus-G (VSV-G) were detected using high throughput imaging or flow cytometry systems. While these approaches were useful, their implementation to different cell types, cargo proteins and various environmental conditions remains difficult.

Indeed, a current challenge is to understand how secretory pathways are specialized and regulated following intrinsic demands, external cues, and eventually altered in diseases. Towards this objective, versatile platforms are needed to uncover factors involved in protein transport in various physiological and pathophysiological contexts. Recent developments of the bacterial Clustered Regulatory Interspaced Palindromic Repeats (CRISPR)-associated nuclease Cas9 technology paired with libraries of single-guide RNAs (sgRNAs), have been successfully used to perform pooled genome-wide screening (Shalem et al., 2015), where targeted genes can be disrupted (Shalem et al., 2014), gene expression inhibited (CRISPRi) or activated (CRISPRa) (Gilbert et al., 2014).

Here, we developed a new and efficient strategy using a pooled genome-wide CRISPRi screen to identify genes involved in protein trafficking and secretion in mammalian cells, and we highlight the contribution of newly identified factors in these processes.

## Results and discussion

### Dual fluorescent reporter for protein transport

We first established a HeLa cell line stably expressing the GFP-tagged TAC protein (interleukin-2 receptor), and the nuclease-dead Cas9 (dCas9) fused to the KRAB repressor protein (HeLa TAC-GFP dCas9-KRAB cells). TAC protein, which contains a single transmembrane domain (TMD) and localizes at the cell surface, represents a straightforward reporter to investigate protein transport along the ER-Golgi secretory pathway (**Fig. 1A to C and Fig. S1A to F**) (Stanley and Lacy, 2010). dCas9-KRAB fusion protein in association to sgRNA targeting the transcription start site (TSS) of specific genes (CRISPRi system) is a powerful tool to inhibit gene expression (Gilbert et al., 2013). We first tested our reporter system by incubating cells with brefeldin A (BFA), an inhibitor of protein transport through the ER and Golgi membranes (Misumi et al., 1986). Flow cytometry analysis demonstrated a decrease of TAC surface expression upon BFA treatment (**Fig. S1G**). Incubation of cells with trypsin, a protease that cleaves most proteins expressed at the cell surface, strongly reduced TAC surface expression that recovered after 18 h in trypsin-free medium. In these conditions, BFA treatment prevented TAC surface expression recovery (**Fig. 1D**).

**Figure 1:**
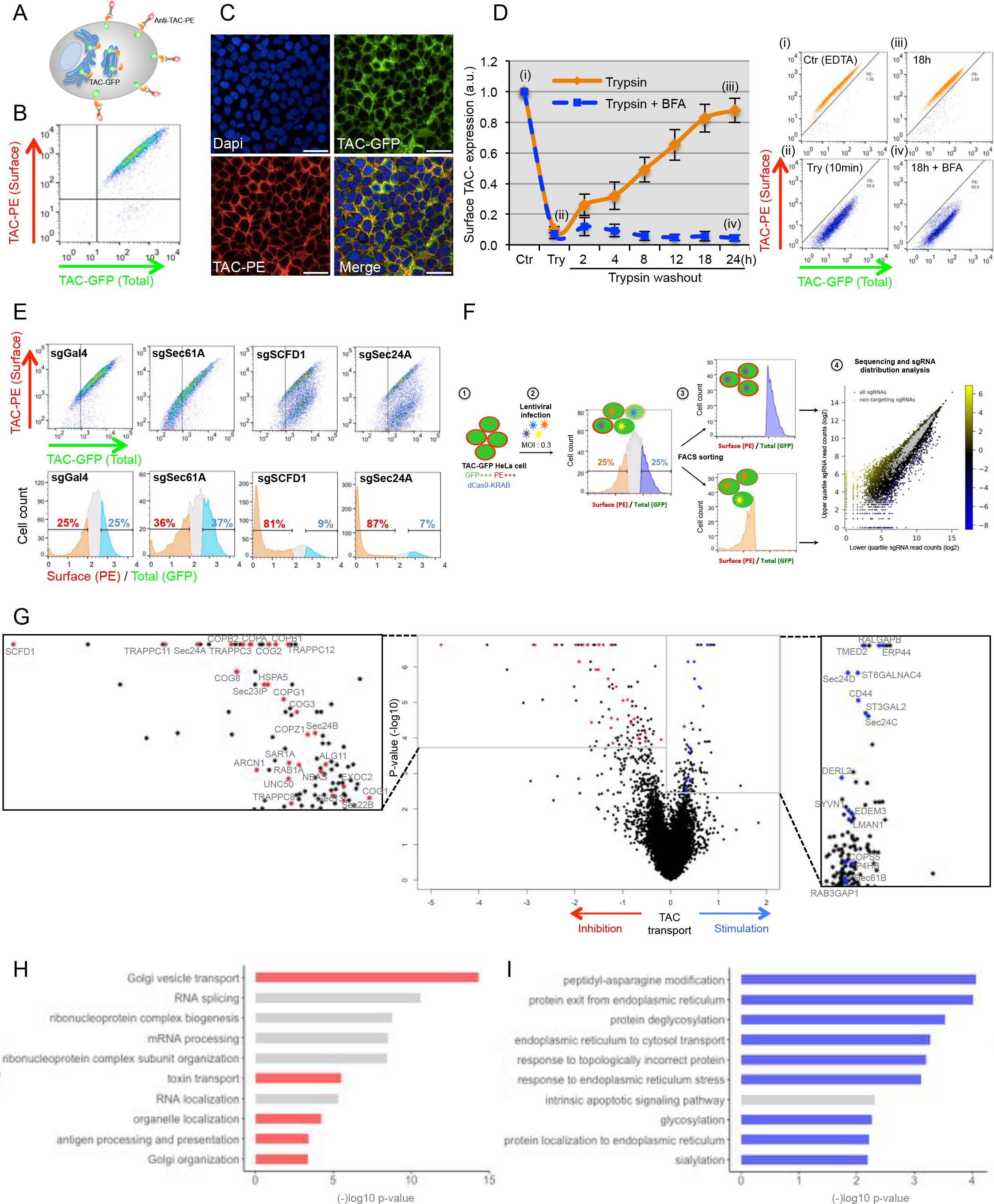
Genome-wide CRISPRi screen for protein transport. (A) Schematic representation of the dual fluorescent reporter. (B and C) Surface and total TAC expression in HeLa TAC-GFP cells were analyzed by flow cytometry (B) and by immunofluorescence microscopy. Scale bar: 50 μm (C). (D) HeLa TAC-GFP cells were detached after incubation with 0.5 mM EDTA (Control (i)) or trypsin (ii) for 10 min and then cells were plated in complete medium in the presence (iv) or absence (iii) of 500 ng/ml BFA. At indicated times, surface and total TAC expression were analyzed by flow cytometry. Surface TAC expression was calculated as the ratio of the mean of fluorescence of surface and total TAC expression (mean ± s.d., n=3). (E) In control HeLa TAC-GFP dCas9-KRAB cells or in cells depleted for Sec61A, SCFD1 and Sec24A, surface and total TAC expression were analyzed by flow cytometry. (F) Schematic representation of the pooled CRISPRi screening workflow. HeLa TAC-GFP dCas9-KRAB cells (1) were infected with the CRISPRi-v2 library (2). Then, using flow cytometry based sorting, two cell fractions representing a low and high surface/total TAC expression ratio were collected (3), and the abundance of each sgRNA in the two fractions was assessed by sequencing (4). (G) Volcano plot showing for each gene, the knockdown effect on TAC transport and P-value (−log10) of phenotype. Screen replicates were averaged. (H and I) Gene Ontology term enrichment analysis among the top 100 genes whose depletion results in (H) TAC transport inhibition, and (I) stimulation.

We then validated our cell line by analyzing TAC expression after knockdown of genes essential for secretory pathway function. Cells were infected with lentivirus to express control sgRNA (Gal4) or sgRNAs targeting the TSS of SCFD1, which regulates SNARE complex assembly (Carr and Rizo, 2010), Sec24A, a COPII vesicular coat subunit (Wendeler et al., 2007), or Sec61A, which promotes translocation of secreted and membrane proteins into the ER (Voorhees and Hegde, 2016). Tight quality-control is associated with the co-translational insertion of nascent polypeptide chains upon emergence from the ribosomal exit tunnel, and a translocation defect of nascent polypeptide chains in the ER leads to their degradation (Brandman and Hegde, 2016). For this reason, downregulation of Sec61A is an appropriate control to monitor TAC expression when TAC translation is altered. Seven days after infection, total and surface TAC expressions were monitored by flow cytometry. Sec61A knockdown, as expected, reduced both surface and total TAC expression. However, SCFD1 and Sec24A knockdown strongly reduced TAC surface transport without affecting the total TAC expression (**Fig. 1E and Fig. S1H**). Cells were also analyzed for the ratio surface/total TAC expression. Upon SCFD1 and Sec24A knockdown, a strong enrichment in the lower quartile was observed related to a decrease of TAC surface expression. Interestingly, no enrichment in one or the other quartile was detected in Sec61A-depleted cells, where both surface and total TAC expression were reduced (**Fig. 1E)**. Altogether, these results established our cell line as an efficient and specific cellular platform to monitor secretory pathway function, and prompted us to use it to perform a pooled genome-wide CRISPRi screen.

### Genome-wide CRISPRi screen mostly identifies genes involved in protein transport

HeLa TAC-GFP dCas9-KRAB cells were transduced with the CRISPRi-v2 library (Horlbeck et al., 2016a) at a low MOI (0.3), and selected for stable integration with puromycin. Seven days after transduction, cells were separated by FACS into a high and low surface/total TAC expression ratio, selecting upper and lower quartiles, respectively. Genomic DNA was then isolated and read counts of sgRNAs obtained by deep sequencing to determine the abundance of each sgRNA in the two cell fractions (**Fig. 1F**). Subsequent analysis using MAGeCK (Li et al., 2014) allowed the identification of many genes whose downregulation modified TAC transport at the cell surface (**Table S1, and Fig. 1G**). In the lower quartile, Gene Ontology (GO) term enrichment analysis of the 100 top ranked genes revealed that knockdown of genes encoding components of “Golgi vesicle transport” was the main functional category inhibiting TAC transport (**Fig. 1G and H**). Genes involved in “toxin transport”, “antigen processing and presentation”, “Golgi organization” were also highly enriched. In the upper quartile, the knockdown of genes encoding known components involved in “protein exit form the ER”, “response to topologically incorrect protein”, “ER to cytosol transport”, “response to ER stress” as well as components involved in post-translational modification were the main functional categories promoting TAC transport (**Fig. 1I**). Altogether, these results clearly demonstrated the value of our approach to identify genes involved in secretory pathway function and organization.

### A secondary screen identifies new factors required for protein transport and secretion

Sixty-three genes identified either in the lower or upper quartiles were selected for a secondary screen using a secretion assay based on the detection of HRP fused to a signal sequence (ss-HRP) (Von Blume et al., 2009). Selected genes were knocked down in HeLa-ss-HRP after transfection of smart pool short-interfering RNAs (siRNAs) pre-arrayed in a 96-well plate. After 3 days, cells were washed, incubated with complete medium, and cell lysates and medium harvested after 8 h for HRP quantification. As a positive control, in SCFD1-depleted cells, a strong signal in the cell lysate associated to a low signal detected in the medium demonstrated a clear defect of HRP secretion and its resulting accumulation inside cells (in red on **Fig. 2A**). Interestingly, this approach identified several genes with poorly characterized function, for which the knockdown inhibited or promoted HRP secretion. They included for example WDR7, YPEL5, TMEM167A, FAM46A, as well as the ubiquitin specific peptidase USP32 and the G protein coupled receptor GPR162. Future analyses will help to dissect their role. We selected 4 genes for further analysis (in green on **Fig. 2A**), those encoding the Coiled-Coil Domain-Containing Protein 151 (CCDC151), and 157 (CCDC157), C10orf88, and the Tetratricopeptide Repeat Protein 17 (TTC17). They were highly enriched in the lower quartile of the CRISPRi screen, their knockdown strongly inhibited HRP secretion, and they encode proteins with a poorly defined or unknown function.

**Figure 2:**
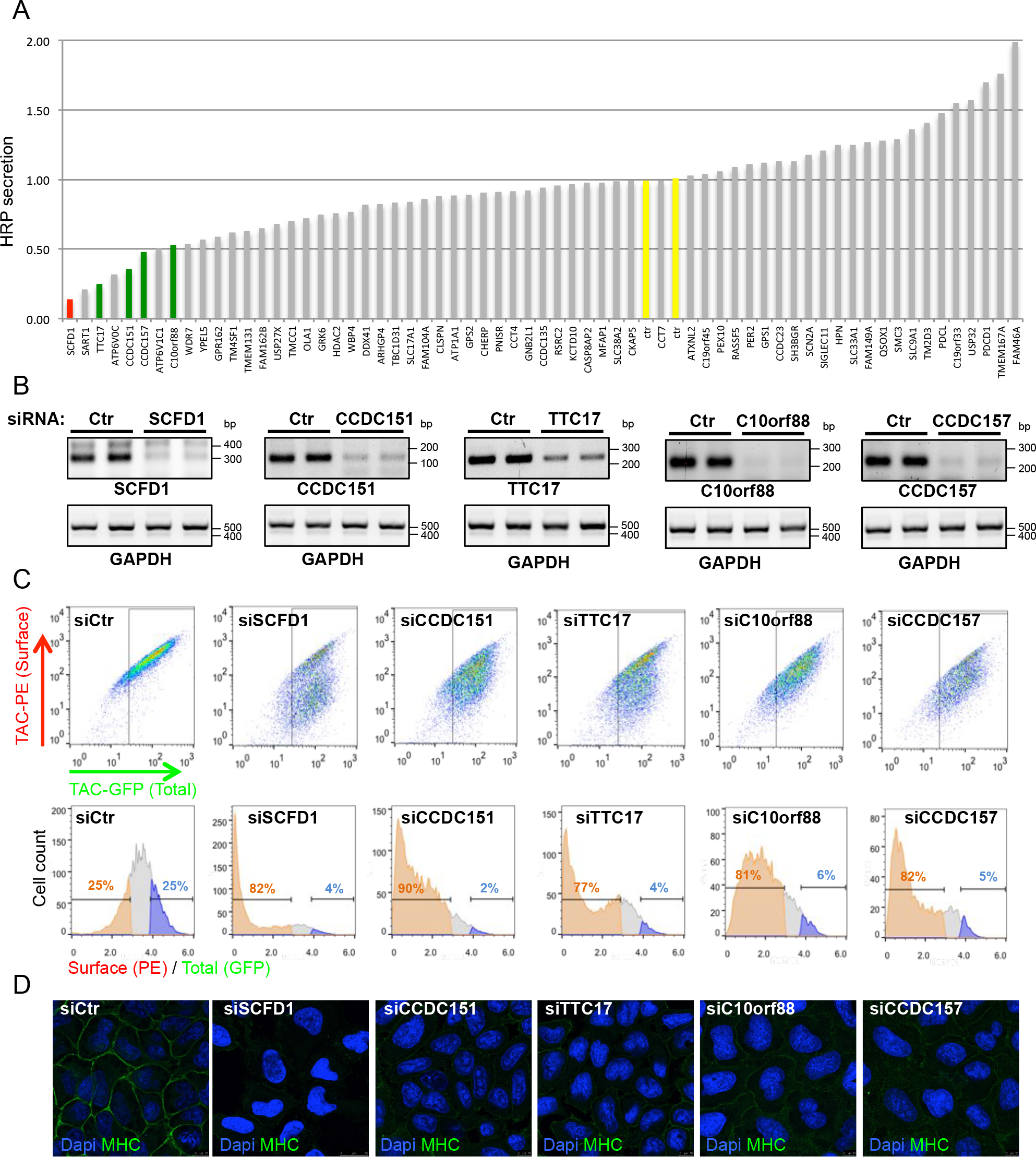
Identification of new factors for protein transport and secretion. (A) HeLa-ss-HRP cells were plated in a 96-well plate pre-arrayed with siRNA. After 3 days, cells were washed and incubated for 8 h with complete medium. HRP secretion was monitored from the supernatant by chemiluminescence and normalized to internal HRP activity (n=2). (B to D) HeLa TAC-GFP cells were transfected with indicated siRNAs and after 3 days, knockdown efficiency was monitored by RT-PCR (B), surface and total TAC expressions were analyzed by flow cytometry (C), and the surface MHC expression was analyzed by immunofluorescence microscopy (D).

First, we aimed to confirm the effect of their knockdown on TAC protein transport. HeLa TAC-GFP cells were transfected with specific siRNA and, after three days, knockdown efficiency monitored by RT-PCR (**Fig. 2B**). Flow cytometry analysis revealed that downregulation of the selected genes inhibited TAC expression at the cell surface, confirming that they were true hits (**Fig. 2C**). We also found that their siRNA-mediated downregulation reduced the transport of the Major Histocompatibility Complex-I (MHC-I), an endogenous cargo protein trafficked along the ER-Golgi secretory pathway to the cell surface (**Fig. 2D**). Altogether, these results demonstrated that the selected candidate genes were required for both exogenous and endogenous cargo protein transport.

### The newly identified factors are essential for secretory pathway organization

Next, we assessed the role of newly identified factors on the morphology of ER, ER exit site (ERES), *cis* Golgi membranes and *trans* Golgi network (TGN) using antibodies targeting calnexin (CNX), Sec31A, GM130 and TGN46, respectively. As a control, in SCFD1-depleted cells, Golgi membranes were completely fragmented, characterized by GM130- and TGN46-positive punctae dispersed throughout the cytoplasm. Interestingly, this phenotype was also observed after CCDC151 and C10orf88 downregulation (**Fig. 3A and C and** **Fig. S2A**). Conversely, after TTC17 and CCDC157 knockdown, Golgi membranes remained localized in the perinuclear area, but presented an extensively dilated and/or expanded organization and a very specific ring-shaped structure, respectively (**Fig. 3D and E and Fig. S2A**). ERES distribution was only slightly altered upon candidate gene knockdown (**Fig. 3B**), and no obvious changes were observed in the distribution and morphology of the ER (**Fig. S2B**).

**Figure 3:**
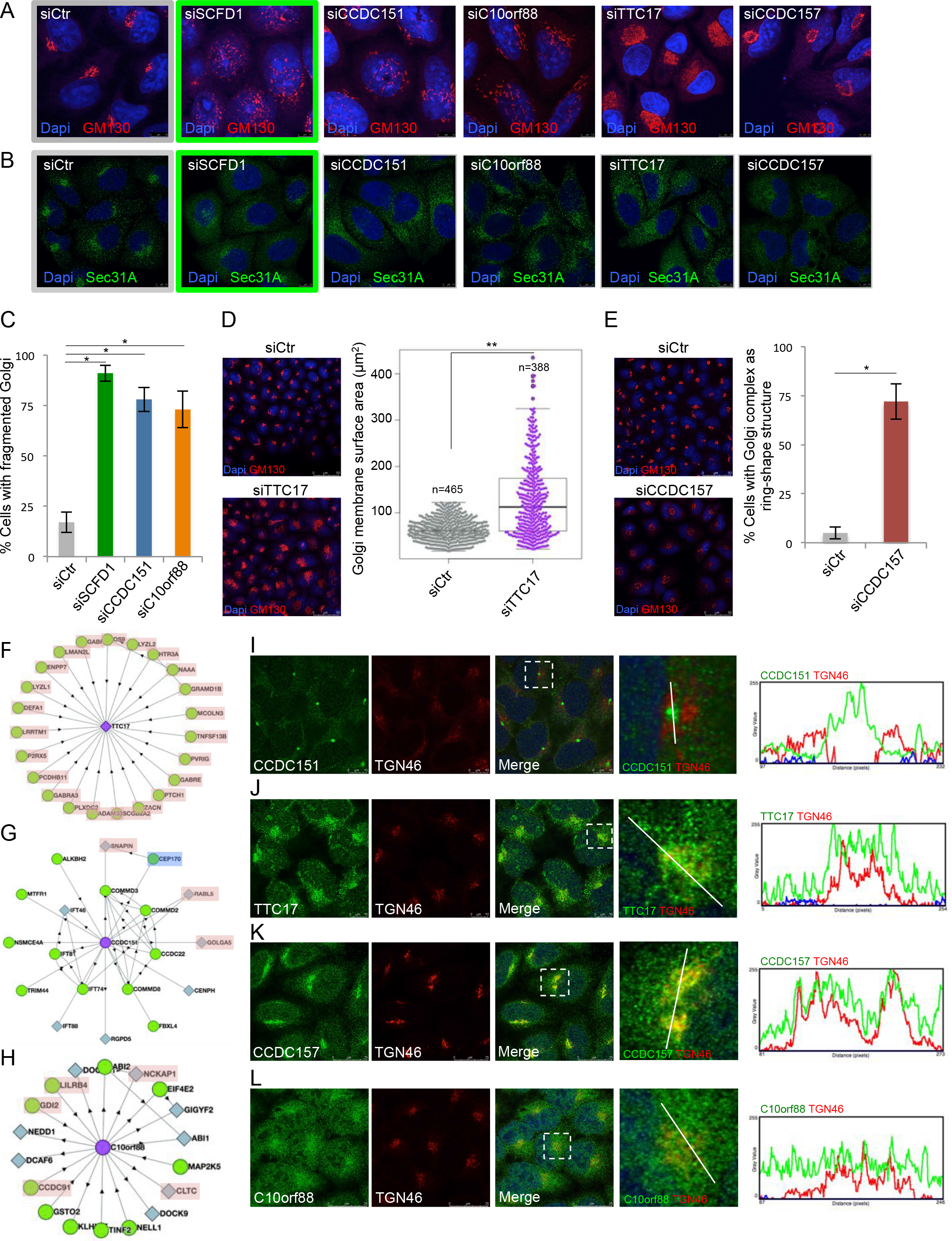
Newly identified factors are essential for secretory pathway organization. (A and B) HeLa cells were transfected with indicated siRNAs and after 3 days, analyzed by immunofluorescence microscopy using antibodies against GM130 (A) and Sec31A (B), respectively. (C) Quantification of the percentage of cells with fragmented Golgi complex after transfection with indicated siRNAs. Cells (200) were counted (mean ± s.d., n=3). (D and E) Left panel. HeLa cells were transfected with indicated siRNAs and after 3 days, analyzed by immunofluorescence microscopy using an anti-GM130 antibody. Right panel. (D) Quantification of Golgi surface area. Control cells and TTC7-depleted cells were analyzed using CellProfiler (**P<0.01). (E) Quantification of the percentage of cells with Golgi membranes with a ring-shape structure. Cells (200) were counted (mean ± s.d., n=3). (F to H) BioPlex analysis displaying interacting partners for (F) TTC17, (G) CCDC151 and (H) C10orf88. Functional proteins acting along the secretory pathway or cargo proteins transported along the ER-Golgi apparatus are highlighted in red. (I to L) Left panels. HeLa cells were processed for cytosolic washout. Then, the co-localization of CCDC151 (I), TTC17 (J), CCDC157 (K) and C10orf88 (L) (green) with TGN46 (red) was assessed by immunofluorescence microscopy. Right panel. Intensity profile graphs of co-localization for the indicated protein with TGN46.

To address the role of the selected factors, we looked for interacting partners. BioPlex network encompasses results aiming at identifying binding partners of proteins encoded by the human genome (Huttlin et al., 2017, 2015). Interrogation of this resource identified TTC17-interacting proteins including exclusively ER and Golgi resident proteins and cargo proteins transported along the secretory pathway (**Fig. 3F**). Interacting partners of CCDC151 include the SNARE-associated protein SNAPIN, GTPase Rab-Like protein 5 (RABL5) and Golgin A5 (**Fig. 3G**). Interacting partners of C10orf88 include Rab protein GDI2, clathrin Heavy chain, clathrin adaptors accessory protein CCDC91 and plasma membrane proteins NCKAP1 and LILRB4 (**Fig. 3H**). The BioPlex resource does not contain a network for CCDC157. Taken together, these data support a direct role of the selected factors on the secretory pathway function, prompting us to uncover their site of action.

A Pfam domain analysis (Finn et al., 2016) or TMHMM software (Krogh et al., 2001) did not predict TMD for the selected factors, whereas HMMTOP algorithm (Tusnády and Simon, 2001) yielded 1 TMD for TTC17. Moreover, their subcellular distribution has not been unambiguously established. To assess localization of the selected factors, we performed immunofluorescence microscopy on HeLa cells, after permeabilization with digitonin and extensive washing, using specific antibodies directed against the endogenous proteins. This approach revealed a close proximity between CCDC151 and the centrosomal marker Centrin-3 (**Fig. 3I and Fig. S2C**). This location is consistent with the interacting partners of CCDC151 revealed by BioPlex and the Golgi membrane phenotype observed in CCDC151-depleted cells (**Fig. 3A and C**). Indeed, interacting partners of CCDC151 included CEP170 (**Fig. 3G**), a key centrosome component that functions as a microtubule-organizing center, essential for the perinuclear localization of Golgi membranes (Rios and Bornens, 2003). Hence, we hypothesized that CCDC151 is a bridging factor between the centrosome and Golgi membranes via interaction with the Golgin A5 and CEP170, thereby contributing to the maintenance of Golgi architecture. Disruption of this functional link may result in Golgi membrane dispersion throughout the cytoplasm after depletion of CCDC151 (**Fig. 3A**). Of interest, our results also revealed the presence of TTC17 and CCDC157 on Golgi membranes (**Fig. 3J and K**). A residual signal of C10orf88 was mainly diffuse in the cytoplasm (**Fig. 3L**). The subcellular location of TTC17 and CCDC157, and the consequence of their knockdown on Golgi membranes prompted us to further characterize these two factors.

### CCDC157 is required for fusion events with Golgi membranes

A permanent flux of membranes goes to and from the Golgi complex. BFA has been regularly used to study Golgi membrane dynamics, inducing a complete fusion of Golgi membranes with the ER and endosomes via tubule formation (Lippincott-Schwartz et al., 1989) and the relocation of peripheral matrix Golgi proteins such as GM130 and GRASP65 to the ERES (Mardones et al., 2006). First, we tested whether Golgi membrane reorganization was altered upon BFA treatment in TTC17- and CCDC157-depleted cells. HeLa cells were transfected with specific siRNAs and after 3 days, treated with BFA. Immunofluorescence microscopy revealed that GM130 completely dispersed upon 30 min of BFA treatment in control cells and in most of CCDC157- and TTC17-depleted cells (**Fig. 4A**). Next, we tested the effect of CCDC157 and TTC17 knockdown on Golgi membrane reassembly after BFA washout, which is mediated by fusion events. In control and TTC17-depleted cells, Golgi membrane reassembly occurred in most cells after 60 min, while in CCDC157-depleted cells, GM130 remained completely dispersed (**Fig. 4A**). These results suggested that Golgi membrane reassembly after BFA washout was strongly dependent on CCDC157.

**Figure 4:**
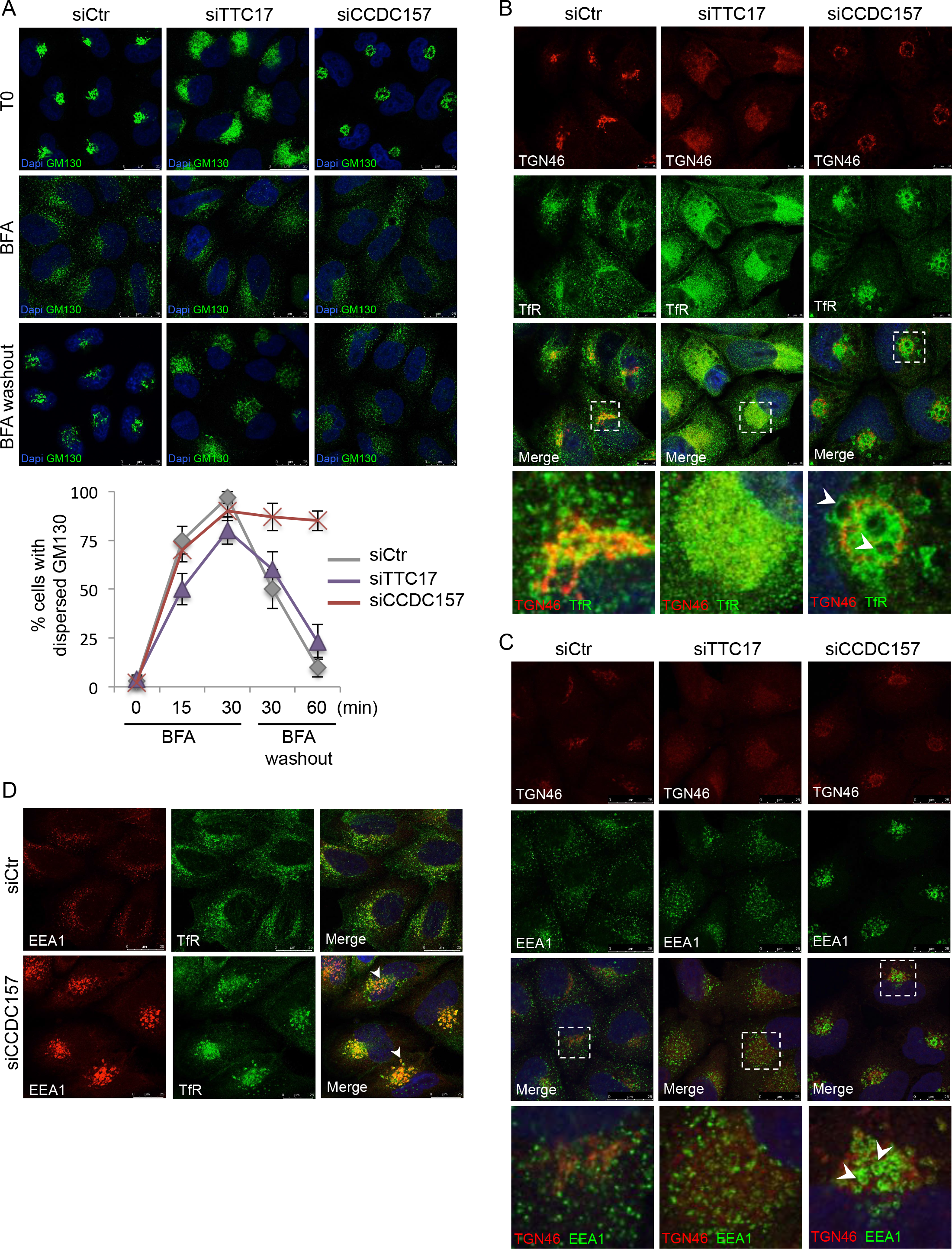
CCDC157 is required for fusion events with Golgi membranes. (A) Control HeLa cells or cells depleted for TTC17 and CCDC157 were incubated with 5 μg/ml BFA and at indicated times, cells were analyzed by immunofluorescence microscopy using an anti-GM130 antibody. For BFA washout, cells were incubated with BFA for 30 min, washed extensively and incubated in BFA-free medium for 30 or 60 min before immunofluorescence microscopy analysis. For quantification, 200 cells were counted for each condition (mean ± s.d., n=3). (B and C) Control HeLa cells or cells depleted for CCDC157 and TTC17 were analyzed by immunofluorescence microscopy using anti-TGN46 and anti-TfR antibodies (B), and anti-TGN46 and anti-EEA1 antibodies (C). Arrowheads point at enlarged TfR- and EEA1-positive vesicles, surrounding by Golgi membranes. (D) Immunofluorescence microscopy analysis using anti-TfR and anti-EEA1 antibodies in control cells and CCDC157-depleted cells.

We next assessed the impact of CCDC157 knockdown on the distribution of transport carriers known to fuse with Golgi membranes. We performed immunofluorescence microscopy experiments using antibodies targeting the transferrin receptor (TfR), which is constitutively transported from the TGN to the cell surface and retrieved to Golgi membranes after endocytosis (Snider and Rogers, 1985), and EEA1, a marker of early endosomes, critical for endosomal trafficking (Barysch et al., 2009). In TTC17-depleted cells, the distribution of TfR and EEA1 were not markedly affected compared to control cells (**Fig. 4B and C**). However, in CCDC157-depleted cells, the distribution of both TfR and EEA1 were strongly altered, most of the signal being restricted to the perinuclear area in the form of enlarged TfR- and EEA1-positive vesicles, surrounded by Golgi membranes (**Fig. 4B to D**). These results suggested that, in the absence of CCDC157, incoming carriers deriving from plasma membrane and endosomes accumulate and collapse in close proximity of Golgi membranes, strengthening CCDC157 as an important factor for the fusion of transport carriers with the Golgi complex.

### TTC17 is required for the polarized arrangement of Golgi cisternae and post-translational modifications

In addition to being a gate entrance and a major sorting station, the Golgi apparatus allows post-translational modifications such as glycosylation. This process is spatially regulated, being dependent on accurate localization of specific enzymes acting sequentially from the *cis*- to *trans*- Golgi cisternae. Does altered Golgi membrane organization in CCDC157- and TTC17-depleted cells impact Golgi polarization and create glycosylation defects? To address this, we first assessed the localization of GM130 and TGN46, *cis*- and *trans*-Golgi markers respectively, in cells incubated in presence or absence of nocodazole. Nocodazole is a microtubule-depolymerizing agent that causes Golgi stack dissociation from each other and their dispersion (Cole et al., 1996), facilitating the visualization of Golgi-resident protein segregation in individual Golgi stacks (**Fig. 5A and D**). In control cells, GM130 and TGN46 staining only partially co-localized. Similar results were obtained in CCDC157-depleted cells. Conversely, in TTC17-depleted cells, GM130 and TGN46 staining showed a high degree of co-localization (**Fig. 5A to E**), suggesting that protein localization in their respective cisternae was perturbed with no further segregation of *cis*- and *trans*- Golgi markers. In addition, we observed that upon TTC17 knockdown, nocodazole failed to induce complete Golgi membrane fragmentation although the microtubule cytoskeleton was entirely depolymerized (**Fig. 5C**). We hypothesized that the dramatic alteration of Golgi membrane architecture upon TTC17 depletion prevents efficient Golgi stack dissociation.

**Figure 5:**
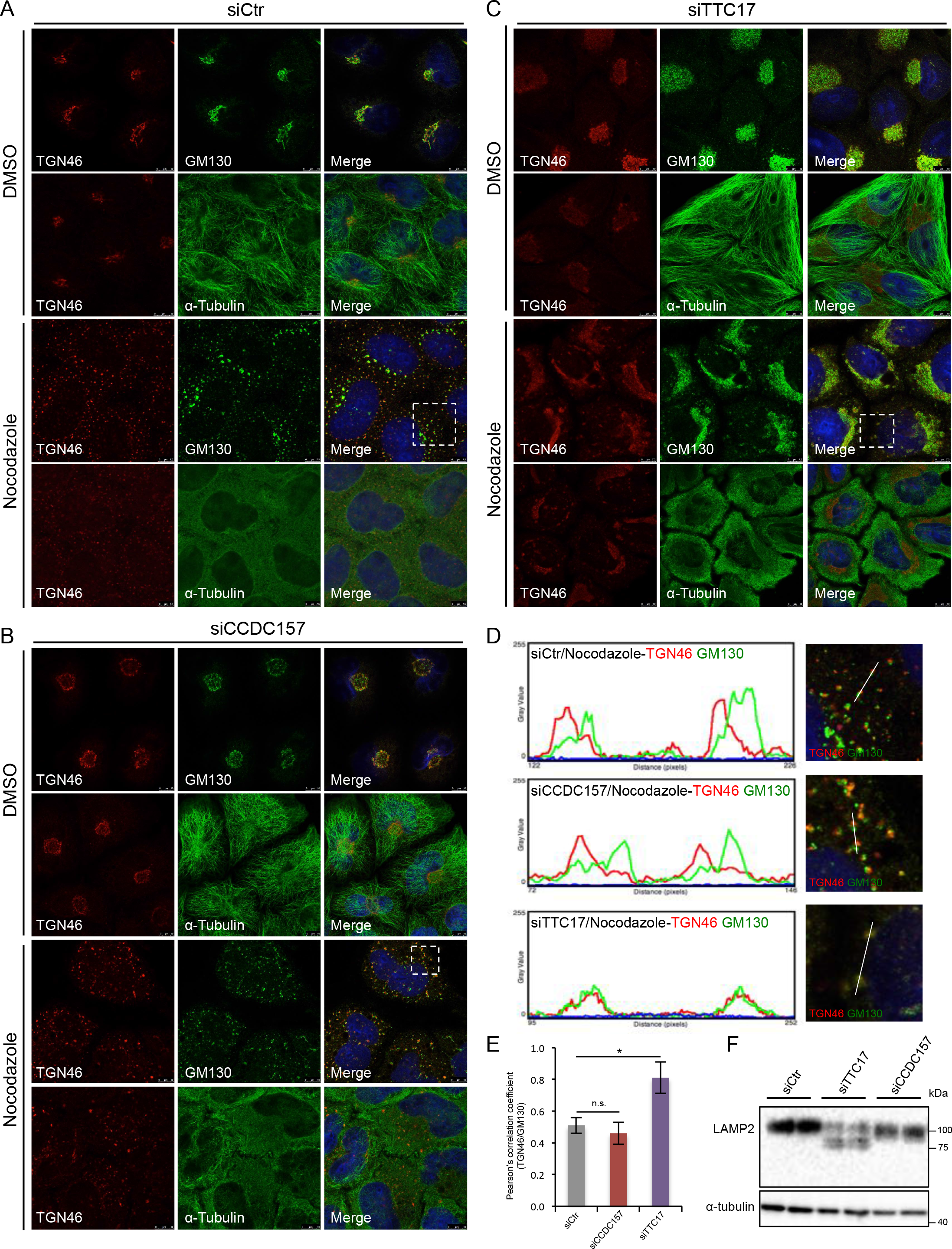
TTC17 is required for the polarized arrangement of Golgi cisternae and post-translational modifications. (A to C) Control HeLa cells (A) or cells depleted for CCDC157 (B) and TTC17 (C) were incubated in the presence or absence of 500 ng/ml nocodazole for 2 h, and analyzed by immunofluorescence microscopy with anti-TGN46 and anti-GM130 antibodies. Depolymerization of the microtubule cytoskeleton was monitored with an anti-α-tubulin antibody. (D) Right panel. Insets from (A), (B) and (C), showing Golgi stack dispersion after nocodazole treatment. Left panel. Intensity profile graphs of TGN46 and GM130 co-localization derived from confocal micrographs shown on the right panel. (E) Pearson’s correlation coefficient of co-localization for TGN46 and GM130 (r) derived from confocal micrographs acquired for control cells (r = 0.51) and cells depleted for CCDC157 (r = 0.46) and TTC17 (r = 0.81) after nocodazole treatment (mean ± s.d., n=8). (F) Total cell lysates of control HeLa cells and cells depleted for TTC17 and CCDC157 were analyzed by immunoblotting with anti-LAMP2 and anti-α-tubulin antibodies.

We next tested whether the loss of Golgi polarization in TTC17-depleted cells impairs cargo protein glycosylation. The lysosomal protein LAMP2 is highly glycosylated during its transport across the secretory pathway, increasing its molecular weight (Stewart et al., 2017). In control and CCDC157-depleted cells, immunoblotting analysis revealed LAMP2 mainly as a fully processed 100 kDa band. However, in TTC17-depleted cells, a smaller form of ~80 kDa became apparent, indicating incomplete LAMP2 processing, likely related to a glycosylation defect (**Fig. 5F**). These results established TTC17 as an important factor for Golgi membrane structure and polarized arrangement of Golgi cisternae, conditioning efficient cargo protein processing and transport.

In summary, we present a powerful unbiased pooled genome-wide CRISPRi screen to identify new factors involved in protein transport and secretion. The ability of our strategy to highlight factors in these processes relied on a combination of key parameters enabling to specifically detect TAC transport alterations and preventing cell death upon systematic gene downregulation. They included an efficient dual fluorescent reporter associated to a CRISPRi-mediated knockdown and an early FACS-based cell selection after lentivirus transduction.

We further characterized two of the newly identified factors. While additional studies will be required to fully decipher their functions, our data suggest that like other coiled-coil proteins (Wong and Munro, 2014; Cheung and Pfeffer, 2016), CCDC157 could be part of a tethering complex required for fusion event with Golgi membranes. On the other hand, phenotypes observed in TTC17-depleted cells associated to its interacting partners involved in sphingomyelin metabolism (ENPP7 and NAAA, **Fig. 3F**) (Tsuboi et al., 2007; Duan et al., 2003), suggest that TTC17 could play a role in the production and distribution of particular lipids, a critical parameter for Golgi complex organization and function (Campelo et al., 2017; van Galen et al., 2014).

Altogether, we anticipate that the list of identified genes as well as the adaptation of this screening platform to specific cargo protein, cell types, or particular environmental challenges can open stimulating perspectives for a deeper understanding of secretory pathway processes in health and disease.

## Materials and methods

### Antibodies, reagents and plasmids

Antibodies used in this study included: mouse monoclonal antibodies anti-TAC conjugated to PE (BioLegend, #356103), anti-GM130 (BD Biosciences, #610822), anti-Sec31A (BD Biosciences, #612351), anti-MHC-I (ThermoFisher, #MA1-70111), anti-EEA1 (BD Biosciences, #610457), anti-alpha-tubulin (ThermoFisher, #62204), anti-LAMP2 (BioLegend, #354302) and anti-Centrin3 (Novus Biologicals, #H00001070-M01), a mouse polyclonal antibody anti-C10orf88 (Abcam, #ab169106), a sheep polyclonal antibody anti-TGN46 (BioRad, #AHP500), rabbit polyclonal antibodies anti-calnexin (Abcam, #ab22595), anti-CCDC151 (Abcam, #ab151469), anti-TTC17 (Atlas antibodies, #HPA038508), anti-CCDC157 (GeneTex, #GTX45090), anti-transferrin receptor (Abcam, #ab84036), secondary antibodies donkey anti-sheep IgG-Alexa Fluor 568 (#A21099), goat anti-mouse IgG-Alexa Fluor 488 (#A11001), goat anti-mouse IgG-568 (#A11004), goat anti-rabbit IgG-568 (#A11011), goat anti-rabbit IgG-488 (#A11034) from ThermoFisher and a secondary antibody mouse IgGκ conjugated to HRP (Santa Cruz Biotechnology, #sc-516102).

Reagents used in this study were obtained from the following sources. Dulbecco’s phosphate buffered saline ((D-PBS, #D8662), BFA (#B6542), nocodazole (#M1404), puromycin (#P9620), G-418 (#4727878001), protease inhibitor cocktail (#05892953001), trypsine (#T3924), penicillin/streptomycin (#P0781), L-glutamine (#G7513) were from Sigma. Q5 HF DNA polymerase (#M0491L), Phusion HF DNA polymerase (#M0530L), Phusion HF reaction Buffer (#B0518S), dNTPs (#N0447S) were from New England Biolabs. T4 DNA ligase (#EL0011), fetal bovine serum (#10270-106), Lipofectamine^®^2000 transfection reagent (#12566014), DH5α bacteria (#18258012), Restore Plus (#46430), ProLong Gold antifade reagent with Dapi (#P36935), OptiMEM (#319885062) were from ThermoFisher. *Trans*IT^®^-2020 transfection reagent (#5405), *Trans*IT^®^-293 transfection reagent (#MIR2700) were from Mirus. Polybrene (#TR-1003-G) and digitonin (#11024-24-1) were from Merck. 0.45μm low protein binding membrane (#28145-479) was from VWR International. NucleoSpin Blood L kit (#740569.10) was from Macherey-Nagel. ECL Western Blotting Detection Reagent (#RPN2106) was from GE Healthcare. Dulbecco’s modified Eagle’s medium ((DMEM), #D6546) was from Molecular Probes. SPRI AMPure XL beads were from Beckman Coulter. White CulturPlate 96-well plates (#6005680) were from PerkinElmer. RNAeasy Plus Mini kit (#74134) was from QIAGEN. Cloned AMV First-Strand cDNA Synthesis kit (#12328032) was from Invitrogen. Mouse IgGκ Isotype (#400111) was from BioLegend. Universal mycoplasma detection kit (#ATCC^®^30-1012K™) was from ATCC.

Plasmids included pCMV6-AC-IL2R-GFP (Origene plasmid #RG215768). pHR-SFFV-dCas9-BFP-KRAB, a gift from Stanley Qi & Jonathan Weissman (Addgene plasmid #46911, (Gilbert et al., 2013)). pU6-sgRNA EF1Alpha-puro-T2A-BFP, a gift from Jonathan Weissman (Addgene plasmid #60955, (Gilbert et al., 2014)). pCMV-VSV-G, a gift from Bob Weinberg (Addgene plasmid #8454, (Stewart et al., 2003)). psPAX2, a gift from Didier Trono (Addgene plasmid #12260). hCRISPRi-v2 genome-wide library, a gift from Jonathan Weissman (Addgene pooled library #83969, (Horlbeck et al., 2016a)).

### Cell culture

All cell lines, HeLa cells (ATCC), HeLa-ss-HPR cells (a gift from Vivek Malhotra), HeLa TAC-GFP cells (this paper), HeLa TAC-GFP dCas9-KRAB cells (this paper) and HEK293T cells (ATCC) were grown in complete medium consisting of Dulbecco’s modified Eagle’s medium (DMEM, Molecular Probes, #D6546) containing 10% fetal bovine serum (FBS, ThermoFisher, #10270-106), 100U/ml penicillin, 100 μg/ml streptomycin (Sigma, #P0781) and 2 mM L-glutamine (Sigma, #G7513) at 37°C with 5% CO_2_. All cell lines were tested every month using ATCC universal mycoplasma detection kit (#ATCC^®^30-1012K™) to confirm they were clear of contamination by mycoplasma.

To generate a HeLa cell line stably expressing GFP-tagged TAC protein, We transfected HeLa cells with pCMV6-AC-IL2R-GFP (Origene plasmid #RG215768) using *Trans*IT^®^-2020 transfection reagent (Mirus, #5405), following the manufacturer’s recommendations, and after two days, transfected cells were selected with 600 μg/ml G-418 (Sigma, #4727878001) during one week. Then, GFP-positive cells were isolated by fluorescence-activated cell sorting (FACS), and single cells were collected in 96-well plates. After expansion into 6-well format, TAC-GFP expression was assessed from clonal lines by flow cytometry and immunofluorescence microscopy analysis. The GFP signal allows monitoring the total TAC expression whereas the pool expressed at the cell surface can be assessed using a phycoerythrin (PE)-conjugated antibody, which recognizes the extracellular domain of the TAC protein. To generate HeLa TAC-GFP dCas9-KRAB cells, we transduced HeLa TAC-GFP cells with lentivirus to express dCas9-KRAB fusion protein (see section “Lentivirus production and transduction”). After two days, transduced cells were sorted for BFP-positive cells.

### siRNA-mediated knockdown

siRNA and smart pools siRNA were transfected in HeLa cells, HeLa-ss-HPR cells, HeLa TAC-GFP cells and HeLa TAC-GFP dCas9-KRAB cells using Lipofectamine^®^2000 transfection reagent (ThermoFisher, #12566014) following the manufacturer’s recommendations. siRNA and smart pools siRNA were purchased from Dharmacon. The siRNA sequences used were: control 5’-CGUACGCGGAAUACUUCGA-3’, SCFD1 5’-AGACUUAUUGAUCUCCAUA-3’, TTC17 5’-GAAUACGGGUGCUGAAGAA-3’, 5’-GGAGAGAGUUAAUCUUUCU-3’, 5’-GGACCUGGAUCUAUAUGAU-3’, 5’-GGACCAGCCUGUACGCUAU-3’, CCDC157 5’-GGAAAGACCUGACGCGCCU-3’, 5’-ACUACGGACCUGCGGCUAA-3’, 5’-CUGAGUGGGAGCACGACAA-3’, 5’-CAGCGUGGAAUCCCAGAUA-3’, CCDC151 5’-GCUCAAGGCCUAUCUAAUG-3’, 5’-GAAGGCUGCCCGAAUACAA-3’, 5’-CGGGAACGCUACAUAAGUG-3’, 5’-GCACAUUACCAGCGUGUAC-3’, C10orf88 5’-GCACAUUGAUGAUAAGAUU-3’, 5’-GAAAUAAGACCGAGUGUCA-3’, 5’-GCUCAGCAGUUGAUGGAUA-3’, 5’-GGGAUACCUCUAAGACAUU-3’. Following incubation for 72 h, total RNA was extracted and knockdown efficiency was assessed by RT-PCR.

### CRISPRi-mediated knockdown

Gene-specific CRISPRi sgRNA oligonucleotide sequences used were: Gal4 5’-GAACGACTAGTTAGGCGTGTA-3’, Sec61A 5’-GGCTAGCACTGACGTGTCTCT-3’, SCFD1 5’-GAGCAGCCAGTATTCGGGAA-3’, Sec24A 5’-GACATGATGACTGGGTTGGA-3’. Forward sgRNA oligonucleotides were designed with 5’ TTG and 3’ GTTTAAGAGC overhangs and reverse sgRNA oligonucleotides were designed with 5’ TTAGCTCTTAAAC and 3’ CAACAAG overhangs. Oligonucleotides were diluted to 100 μM in water and 1 μl of forward and reverse oligo were combined in a 50 μl reaction containing 200 mM potassium acetate, 60 mM HEPES, pH 7.4 and 4 mM magnesium acetate. Oligonucleotides were incubated for 5 min at 95°C and slow cooled (0.1 degrees/sec) for annealing. Annealed oligos were diluted 1/40 and 1 μl of insert was ligated into 10 ng of digested vector (pU6-sgRNA EF1Alpha-puro-T2A-BFP, Addgene plasmid #60955 digested with BstXI and Blpl) using T4 DNA ligase (ThermoFisher, #EL0011) in a final volume of 10 μl. Ligation was allowed to proceed for 1 h. Ligated products were transformed into DH5α bacteria (ThermoFisher, #18258012). After lentivirus production (see section “Lentivirus production and transduction”), TAC-GFP-dCas9-KRAB HeLa cells were infected and knockdown efficiency was monitored 7 days post-transduction by RT-PCR. TAC-GFP expression was assessed by flow cytometry analysis.

### Lentivirus production and transduction

Lentivirus was generated in HEK293T cells using *Trans*IT^®^-293 transfection reagent (Mirus, #MIR2700) following the manufacturer’s recommendations. At 50% confluence, HEK293T cells incubated in complete medium lacking penicillin/streptomycin were transiently transfected with a lentiviral vector, either pHR-SFFV-dCas9-BFP-KRAB (Addgene plasmid #46911) for dCas9-KRAB fusion protein expression, pU6-sgRNA EF1Alpha-puro-T2A-BFP (Addgene plasmid #60955) containing gene-specific CRISPRi sgRNA oligonucleotides, or hCRISPRi-v2 library (Addgene pooled library #83969) for a pooled genome-wide CRISPRi screen, together with packaging plasmids pCMV-VSV-G (Addgene plasmid #8454) and psPAX2 (Addgene plasmid #12260) at a 4:1:3 ratio. Transfection was performed using one 10-cm dish for dCas9-KRAB fusion protein expression and for each specific sgRNA, and using two 15-cm dishes for hCRISPRi-v2 library. After transfection (24 h), the medium was refreshed and after additional 24 h virus was collected and filtered through a 0.45μm low protein binding membrane (VWR International, Radnor, PA, #28145-479). For dCas9-KRAB fusion protein expression and CRISPRi-mediated knockdown, 1 ml of medium was used immediately for infection of ~500,000 target cells plated in 6-well plate in a final volume of 2 ml. For a pooled genome-wide CRISPRi screen, ~200 million TAC-GFP-dCas9-KRAB HeLa cells plated in 30 15-cm dishes were infected at a multiplicity of infection (MOI) of 0.3. Medium was supplemented with 8 μg/ml polybrene (Merck, #TR-1003-G) lacking penicillin/streptomycin. After 24 h, medium was replaced with fresh medium and after an additional 24 h, high BFP-positive cells expressing dCas9-KRAB fusion protein were sorted by FACS. For CRISPRi-mediated knockdown and a pooled genome-wide CRISPRi screen, 48 h after infection, 2μg/ml puromycin (Sigma, St. Louis, MO, #P9620) was added to select transduced cells.

### Pooled genome-wide CRISPRi screen and Illumina library construction and sequencing

Approximately 200 million HeLa cells stably expressing TAC-GFP and dCas9-KRAB fusion protein were transduced with the hCRISPRi-v2 library (Addgene #83969) at an MOI of 0.3 to ensure no more than one viral integration event per cell. The CRISPi-v2 library comprised 104,535 sgRNAs targeting 18,905 human genes. Each sgRNA of the library targeted the TSS of the respective genes in regions of low nucleosome occupancy avoiding genomic DNA compaction to impact Cas9 binding (Horlbeck et al., 2016b; a). Two days after infection, an aliquot of cells was analyzed by FACS to confirm transduction efficiency and then, cells were cultured with 2 μg/ml puromycin (Sigma) to remove uninfected cells. After an 5 additional days, cell were trypsinized, washed extensively and plated with fresh medium for 18 h allowing recovery of transmembrane protein expression at the cell surface. Cells were then detached using 0.5 mM EDTA, resuspended in blocking buffer (OptiMEM containing 5% FBS) and stained for TAC surface expression using a PE-conjugated anti-TAC antibody during 1 h at 4°C. Finally, live cells were sorted by FACS following the PE/GFP fluorescence ratio, selecting the upper and lower quartile, respectively. Ten million cells were sorted for each fraction. Cell sorting was performed using a FACSAria sorter. Two independent screens were performed. After FACS sorting, cells were centrifuged and pellets were frozen until processing.

Genomic DNA was extracted from four distinct cell populations, corresponding to the upper and lower quartile of the PE/GFP fluorescence ratio, from 2 independent screens, using the NucleoSpin Blood L kit (Macherey-Nagel, #740569.10) according to the manufacturer’s instructions. Then, for each sample, all genomic DNA was amplified by PCR to enrich sgRNA sequences. Each 100 μl PCR reaction contained 1x Phusion HF reaction Buffer (NEB, #B0518S), 3% v/v DMSO, 0.4 μM of each forward and reverse primer mix (Integrated DNA Technologies), 0.2 mM dNTPs (NEB, #N0447S), 40 U/ml Phusion HF DNA polymerase (NEB, #M0530L) and 2 μg genomic DNA. PCR primer sequences for the forward (Fw) primers with specific barcode for de-multiplexing were:

5’-aatgatacggcgaccaccgagatctacacgatcggaagagcacacgtctgaactccagtcacCTTGTAgcacaaaaggaaactcaccct-3’
5’-aatgatacggcgaccaccgagatctacacgatcggaagagcacacgtctgaactccagtcacGCCAATgcacaaaaggaaactcaccct-3’
5’-aatgatacggcgaccaccgagatctacacgatcggaagagcacacgtctgaactccagtcacAGTTCCgcacaaaaggaaactcaccct-3’
5’-aatgatacggcgaccaccgagatctacacgatcggaagagcacacgtctgaactccagtcacTAGCTTgcacaaaaggaaactcaccct-3’

The PCR sequence for the common reverse (Rev) primer was: 5’-CAAGCAGAAGACGGCATACGAGATCGACTCGGTGCCACTTTTTC-3’. PCR cycling conditions used included 1x 98°C for 30s; 23x 98°C for 30s, 56°C for 15s, 72°C for 15s, 1x 72°C for 10 min. For each sample, PCR products were pooled, size selected using SPRI AMPure XL beads (Beckman Coulter) following the manufacturer’s recommendations, and quantified on a Qubit and run on a Bioanalyzer (Agilent). The four libraries were pooled to a total concentration of 10 nM and subjected to DNA sequencing on a HiSeq2500 (Illumina) sequencer. Samples were sequenced with the rapid run mode of the single-ended 50 pb, following the manufacturer’s recommendations, using the custom sequencing primer: 5’-GTGTGTTTTGAGACTATAAGTATCCCTTGGAGAACCACCTTGTTG-3’.

For data analysis, raw sequencing reads were aligned to the v2CRISPRi library sequences using Bowtie 2, and data were analyzed using MaGECK algorithm (Li et al., 2014). For genes with multiple independent TSS targeted by the sgRNA library, phenotype and P-values were calculated independently for each TSS. The CRISPRi screen was performed twice.

### Secondary screen using an HRP secretion assay

HeLa cells stably expressing ss-HRP were transfected in 96-well plate pre-arrayed with the indicated smart pools siRNA (Dharmacon) using Lipofectamine^®^2000 transfection reagent (ThermoFisher, #12566014). Briefly, for each well, 3000 HeLa-ss-HRP cells in 80 μl of complete medium lacking penicillin/streptomycin were plated into 96-well plate containing 20 μl of 250 nM siRNA and 0.5% (v/v) lipofectamine 2000 in OptiMEM (ThermoFisher, #319885062). Medium was replaced after 24h. After 2 additional days, cells were washed with medium and incubated with 150 μl of complete medium for 8 h. Then 50 μl of the medium was collected, mixed with ECL reagent (GE Healthcare, #RPN2106) in white CulturPlate 96-well plates (PerkinElmer, #6005680), and the chemoluminescence measured in a Tecan Microplate reader M1000 PRO. For normalization, cells were lysed with lysis buffer supplemented with protease inhibitors to quantify intracellular HRP activity.

### Flow cytometry

After CRISPRi- or siRNA-mediated target gene knockdown, cells were washed with PBS and detached by incubation with 0.5 mM EDTA for 10 min in PBS. Cells were then resuspended in blocking buffer (OptiMEM containing 5% FBS) and live cells were incubated for 1 h at 4°C with an anti-TAC antibody conjugated to PE. GFP and PE fluorescence were collected on a LSR Fortessa (BD Biosciences) and analyzed using FlowJo software (FlowJo, LLC).

### Immunoblot

Cells were lysed for 30 min on ice in lysis buffer (50 mM Tris, pH 7.4, 150 mM NaCl, 0.1% SDS, 1% (v/v) Triton X-100 and 0.5% sodium deoxycholate) supplemented with protease inhibitor coktail (Sigma-Aldrich, #05892953001), 1 mM Na_3_VO_4_ and 25 mM sodium fluoride and centrifuged at 16,000xg for 15 min. Samples were incubated with 1X SDS sample buffer at 95°C for 10 min, resolved by 12% SDS-PAGE and transferred to methanol activated Immobilon^®^-P 0.45 μm PVDF membranes for blotting. Membranes were blocked with 5% (w/v) BSA in PBS containing 0.1% Tween 20 (PBS-T) for 30 min at room temperature, and probed with appropriate primary antibodies overnight at 4°C in PBS-T containing 5% (w/v) BSA. Membranes were then washed 3 × 15 min in PBS-T and incubated with the corresponding HRP-conjugated secondary antibody for 1 h at room temperature in PBS-T containing 5% (w/v) BSA. Membranes were then washed again for 3 × 15 min in PBS-T and signals were detected with ECL Western Blotting Detection Reagent (GE Healthcare, #RPN2106) and acquired with Chemidoc MP Imaging System (Bio-Rad, Hercules, CA). Membranes were stripped with Restore Plus (ThermoFisher Scientific, #46430) according to the manufacturer’s instruction.

### Immunofluorescence microscopy

Cells grown on coverslips were fixed with cold methanol for 10 min at −20°C or 4% (w/v) paraformaldehyde (PFA) in PBS for 15 min at room temperature. Cells fixed with PFA were permeabilized with 0.1% Triton X100 in PBS at room temperature and then incubated with blocking buffer (2.5% (w/v) FCS, 0.1% triton X100 in PBS) for 30 min at room temperature. Cells were then incubated with primary antibody diluted in blocking buffer for 1 h at room temperature followed by PBS wash and incubated with secondary antibody. Secondary antibodies conjugated with Alexa Fluor 488, 568, or 647 were diluted in blocking buffer and incubated 1 h at room temperature. Samples were mounted using ProLong Gold antifade reagent with Dapi (4,6-diamidino-2-phenylindole; ThermoFisher, #P36935). Images were acquired with a Leica SP8 laser confocal scanning microscope with a 40x objective. Co-localization analysis was performed using ImageJ (National Institutes of Health), and the Pearson’s correlation coefficient was calculated using Costes method of thresholding with object Pearson’s analysis using Imaris 8.2.0 by Bitplane AG. Images displayed in figures are representative single Z-slices. For quantification of Golgi membrane area, images were analyzed with the CellProfiler software (Broad Institute).

For cytosol washout prior immunofluorescence microscopy, cells were washed twice with room temperature KHM buffer (125 mM potassium acetate, 25 mM HEPES-KOH, pH 7.2, and 2.5 mM magnesium acetate). Cells were then permeabilized by incubation in KHM buffer containing 30 μg/ml digitonin for 5 min on ice followed by a wash for 5 min at room temperature with KHM buffer. Cells were subsequently fixed with 4% paraformaldehyde and processed for immunofluorescence microscopy. This procedure allows removing the soluble cytoplasmic pool of proteins, highlighting their potential association with intracellular compartments.

### RT-PCR

Total RNA was extracted from HeLa cells using the RNAeasy Plus Mini kit (QIAGEN, #74134) and cDNAs were synthesized with the Cloned AMV First-Strand cDNA Synthesis kit (Invitrogen, #12328032) and oligo d(T)_20_ according to the manufacturer’s instruction. cDNA of indicated genes were amplified by PCR using Q5 HF DNA polymerase (NEB, #M0491L) and the following specific primers: GAPDH Fw primer 5’-ACCACCATGGAGAAGGCTGG-3’, Rev primer 5’- CTCAGTGTAGCCCAGGATGC-3’, PCR product size: 527 bp; SCFD1 Fw primer 5’-GGGGAAGATGAAGGAGCCATA-3’, Rev primer 5’-TCAGCCTCAGAAGGTGCTTG-3’, PCR product size: 319 bp; Sec24A Fw primer 5’-CCATTGTCCTCGTGCATCAT-3’, Rev primer 5’-TGCCAATAAGCCACCTCCTT-3’, PCR product size: 369 bp; Sec61A Fw primer 5’-AGTGGACCTGCCAATCAAGT-3’, Rev primer 5’- CTTTGGCAGAGGAACCTGAG-3’, PCR product size: 379 bp; TTC17 Fw primer 5’-CAGATGACCATGCACGAAAAA-3’, Rev primer 5’-GTTGGCCAAGTTGACAAGAGG-3’, PCR product size: 248 bp; CCDC157 Fw primer 5’-GCAGCCCACCCACTGTAATAA-3’, Rev primer 5’-TGCCCTGGGTGTACTCTCCTA-3’, PCR product size: 236 bp; C10orf88 Fw primer 5’-TACACAACTGCCTGGTGGAGA-3’, Rev primer 5’-CGGTCTTATTTCCCACATGGA-3’, PCR product size: 218 bp; CCDC151 Fw primer 5’-TGAACCAAGAGGCCCTCAAT-3’, Rev primer 5’-GCGCTCGTTCTCCAGTTTCT-3’, PCR product size: 148 bp. The PCR products were resolved on 1.5% agarose gel.

### Statistical analysis

Data are presented as mean ± s.d. Differences in the mean values between 2 groups were assessed by 2-tailed Student *t*-test. *P<0.05 was taken to imply statistical significance. In **figure 3D**, difference between 2 groups was assessed by Mann-Whitney U test. **P<0.01.

### Online supplemental material

**Fig. S1** shows the establishment of a cell line expressing a dual fluorescent reporter for protein transport, the optimization of experimental conditions to assess TAC expression, and the knockdown efficiency after lentivirus infection to express sgRNAs targeting the TSS of Sec61A, SCFD1 and Sec24A. **Fig. S2** shows the effect of gene knockdown on TGN and ER organization, as well as the subcellular localization of CCDC151 at the centrosome. Table S1 shows MaGECK analysis of the CRISPRi screen replicate.

## Acknowledgments

We thank members of the Moreau lab, Malhotra lab and Schekman Lab for helpful discussion. Stephanie J. Popa acknowledges support from a Wellcome Trust PhD studentship. Anupama Ashok acknowledges support from a la Caixa fellowship. Sarah E. Stewart acknowledges support from a BBSRC Future Leader Fellowship (BB/P010911/1). Julien Villeneuve acknowledges support from a Marie Curie international outgoing fellowship within the 7^th^ European community framework program. This work was supported by a Wellcome Trust Strategic Award [100574/Z/12/Z], the Medical Research Council Metabolic Diseases Unit [MRC_MC_UU_12012/5], and the Isaac Newton Trust/Wellcome Trust ISSF/ University of Cambridge research grant.

The authors declare no competing financial interests.

## Author contributions

J. V., K. M., V. M., R. S., J. W. and J. R. conceptualized the experiments. J. V., L. B. and S.J.P. performed and analyzed most of the experiments, with contributions from M. H., A. A., S. E. S., C.M.B., N. B. and K. H. Writing: J. V. and K. M. wrote the original and revised manuscripts with contribution from the other authors.

---

**Figure S1:**
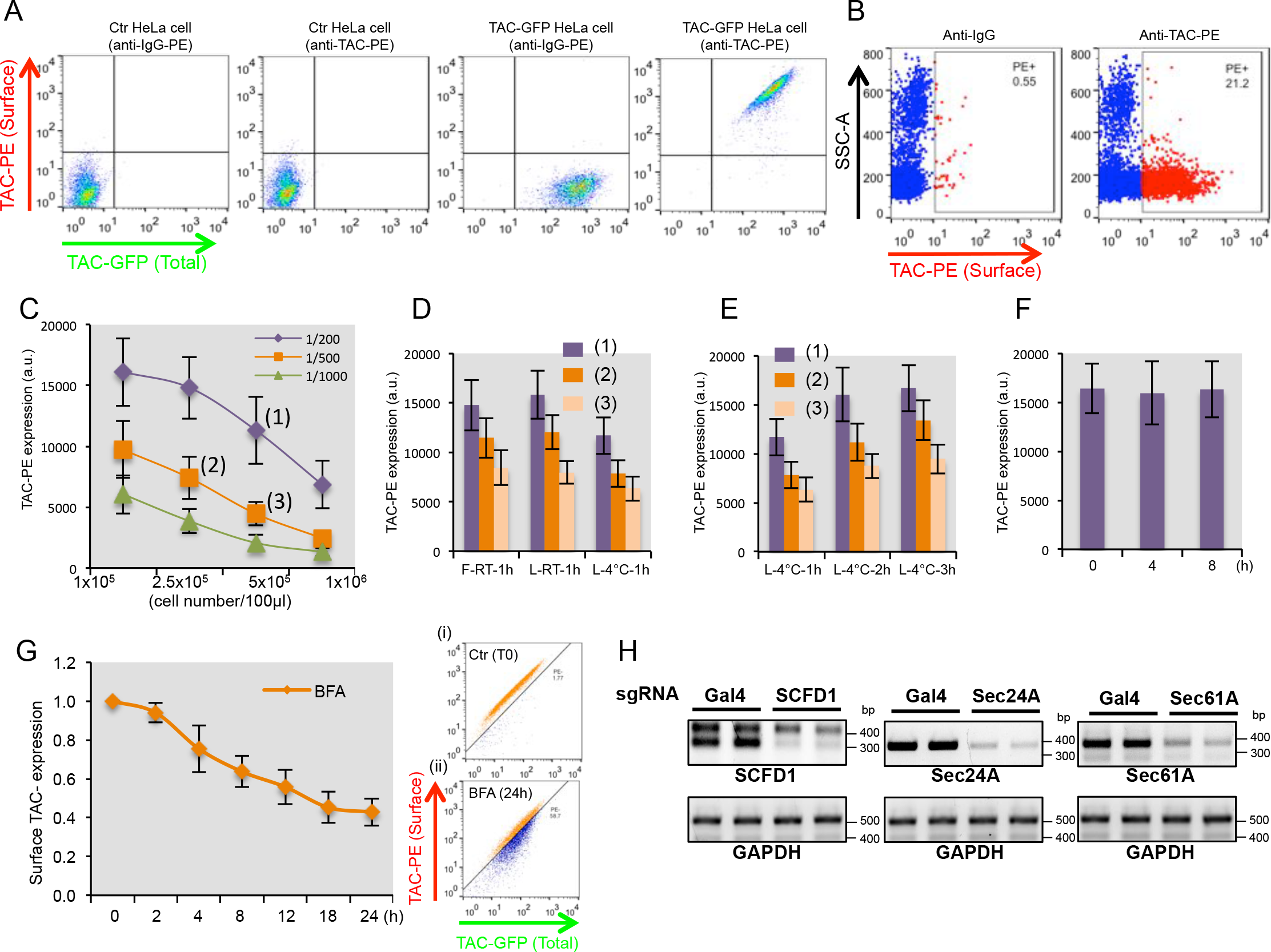
Dual fluorescent reporter for protein transport. (A) Surface and total TAC expression were analyzed by flow cytometry in control HeLa cells and in cells stably expressing TAC-GFP. (B) TAC protein is endogenously expressed on the surface of certain immune cells, such as lymphocytes. To test further the specificity of the anti-TAC antibody used, TAC surface expression was analyzed by flow cytometry in nucleated cells from blood sample. Only a subpopulation of nucleated cells, most likely the lymphocytes expressed TAC proteins. (C to F) Optimization of experimental conditions to monitore TAC expression by flow cytometry. (C) TAC surface expression in HeLa-TAC-GFP cells (from 1×10^5^ to 1×10^6^ cells/100μl) were analyzed by flow cytometry using increasing concentrations of anti-TAC-PE antibody. (D) Using conditions establish in (C), TAC surface expression was analyzed on fixed (F) and Lived (L) cells by incubating cells with the anti-TAC-PE antibody during 1 h at room temperature (RT) or at 4°C. (E) Using conditions establish in (C), TAC surface expression was analyzed on Lived (L) cells by incubating cells with the anti-TAC-PE antibody during 1, 2 or 3 h at 4°C. (F) Lived HeLa-TAC-GFP cells were incubated for 2 h at 4°C with the anti-TAC-PE antibody, followed by flow cytometry analysis at T=0, or after 4 and 8 h. TAC surface expression on lived cells maintain 4°C after staining remain stable even after 8h. This control was essential to ensure the stability of the signal during the step of FACS sorting for the pooled genome-wide CRISPR screen. (G) Left panel. HeLa TAC-GFP cells were incubated in presence or absence of 1 μg/ml BFA and at indicated times, cells were washed with PBS, detached by 10 min incubation in PBS containing 0.5 mM EDTA, and surface and total TAC expression was analyzed by flow cytometry. Surface TAC expression was calculated as the ratio of the mean of fluorescence of surface and total TAC expression. Results are shown as the mean ± SD of 3 independent experiments. Right panel. Flow cytometry analysis of Surface and total TAC expression of HeLa TAC-GFP cells at T0 (i) and after incubation in presence of 1 μg/ml BFA during 24 h (ii). (H) HeLa TAC-GFP dCas9-KRAB cells were infected with lentivirus to express control sgRNA (Gal4) or sgRNAs targeting the TSS of SCFD1, Sec24A and Sec61A. Seven days after infection, the knockdown efficiency was monitored by RT PCR.

**Figure S2:**
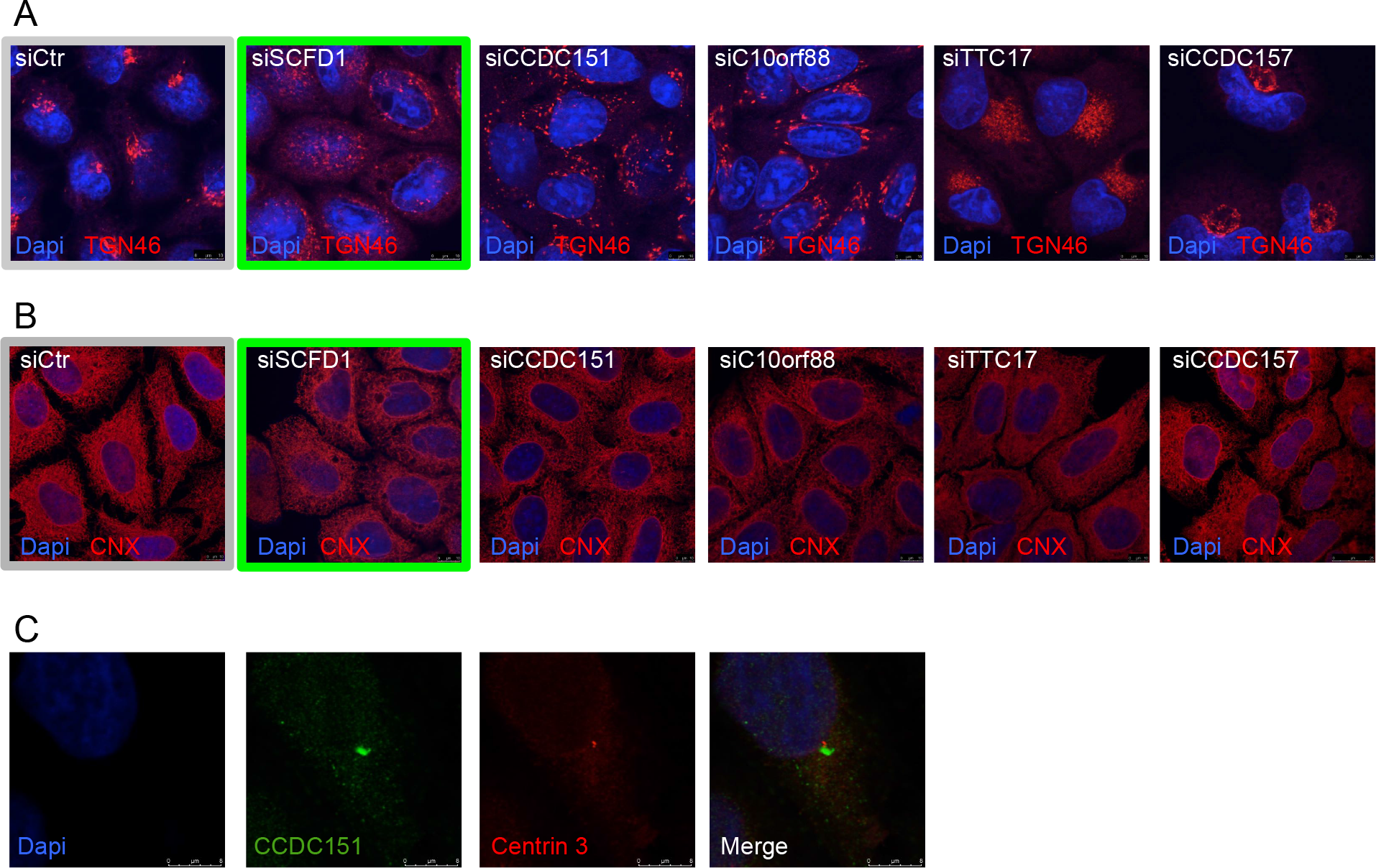
Newly identified factors are essential for Golgi membranes organization. (A and B) HeLa cells were transfected with the indicated siRNA and after 3 days the TGN and the ER organization were assessed by immunofluorescence microscopy using specific antibodies against TGN46 (A) and CNX (B), respectively. (C) CCDC151 localizes at the centrosome. HeLa cells were processed for cytosolic washout. After fixation and permeabilization, the co-localization of the endogenous protein CCDC151 (green) with Centrin 3 (red), a centrosomal marker, was monitored by immunofluorescence microscopy. Nuclei are stained with Dapi.

